# Leveraging genomes and metagenome-assembled genomes unveils novel diversity and transmission between public transport and clinical settings in *Acinetobacter*

**DOI:** 10.64898/2026.09.29.755473

**Authors:** Rafael López-Sánchez, Alejandro Aguilar-Vera, Santiago Castillo-Ramírez

## Abstract

The molecular epidemiology of *Acinetobacter*, which is almost exclusively culture-dependent, has centered mainly on clinical isolates of *A. baumannii*. Yet, this overlooks sources not amenable to microbial culture and many non-*A. baumannii* species. Here, by analysing metagenomic projects from public transport systems worldwide, we show that metagenome-assembled genomes (MAGs) are highly relevant for assessing the unrecognized diversity of *Acinetobacter*. We recovered 839 *Acinetobacter* MAGs, with 269 being high-quality MAGs. The high-quality MAGs were across more than 20 well-defined *Acinetobacter* species; however, five species (*A. nosocomilalis, A. ursingii, A. radioresistens, A. schindleri*, and *A. variabilis*) had the most MAGs. Pangenome and phylogenetic diversity analyses of 685 genomes plus the 269 high-quality MAGs from those 5 species showed that MAGs are an important source of both lineage and gene diversity. Also, F_ST_ analyses and genome phylogenies show that for some species there is no genetic differentiation between MAGs and clinical isolates, implying transmission between hospitals and public transport settings. In addition to finding the antibiotic resistance genes (ARGs) in MAGs, we implemented an *ad hoc* strategy (leveraging the use of genomes) to recover considerably more ARGs. PCoA and PERMANOVA analyses show that, whereas in *A. nosocomialis* and *A. ursingii*, the ARG profiles of MAGs and clinical genomes are different, in *A. radioresistens* and *A. schindleri*, the ARG profiles are not different. Finally, some clinically important ARGs, such as *sul1* and *sul2, aadA*, and *blaOXA*_*23*_, were found in all or some of the 5 species. Our study shows how combining MAGs and regular genomes is a powerful strategy for better appraising the genomic diversity (including the resistome) of *Acinetobacter*. On a broader level, our study showcases the importance of MAGs for genomic surveillance of man-made environments with clear public health implications.

## Introduction

*Acinetobacter* is a bacterial genus very diverse both in terms of sources and metabolic capacity [1]. Nonetheless, the genus is mainly associated with hospital infections and mostly just with one species, namely *Acinetobacter baumannii* [2]. Yet, as of late August 2026, this genus includes 95 species with valid, published names (https://lpsn.dsmz.de//genus/acinetobacter); however, the information on the different species is highly skewed. Whereas for *A. baumannii* there is a lot of information; as of 16 September 2026, there are almost 19,000 publications on PubMed for this species. For other *Acinetobacter* species (such as *A. ursingii* or *A. variabilis*), there is barely any information. For instance, the query “Acinetobacter ursingii” yielded just 53 publications; the query “Acinetobacter schindleri” yielded just 47 publications; while the query “Acinetobacter variabilis” yielded only 10 publications. Thus, for several species within this genus, there is very little knowledge. It is also relevant to mention that, as far as the habitat of the *Acinetobacter* species is concerned, some species have been noted to be able to inhabit both human and non-human sources [3-6].

In terms of understanding the habitat range of the *Acinetobacter* species, metagenomics has great potential in that it bypasses the culture dependence of standard microbial characterization techniques [2, 7]. Thus, metagenomics is very convenient when obtaining cultures from particular sources or hosts is rather difficult or impossible. Indeed, recent studies have shown the usefulness of metagenomics, in general, and metagenome-assembled genomes (MAGs), in particular, to study atypical sources of *Acinetobacter* species. For instance, we recently showed that some unexpected places can have *A. baumannii* lineages, among them the New York subway system [8]. Another study found *A. calcoaceticus* in the microbiota of field-grown tomato plants [9]. Also, MAGs for *A. baumannii* and *A. pittii* were recovered from wound tissue biopsies [10]. Importantly, besides hospitals, we know almost nothing about whether or not *Acinetobacter* species can inhabit other human-made environments. Among these environments, public transport systems are bound to be very relevant in terms of infectious disease dissemination in cities. Thus, here we analyse metagenomic projects from public transport systems to evaluate potentially overlooked diversity of the genus *Acinetobacter*. We pay special attention to the transmission of isolates between clinical and public transport settings as well as the resistome of the most prevalent *Acinetobacter* species dwelling in public transport systems.

## Results

### *Acinetobacter* inhabits the public transport system

To explore whether *Acinetobacter* species dwell in public transport settings, we analysed metagenomic data from public transport systems. We analysed publicly available metagenomic sequencing data from 15 BioProjects (see Figure 1A) deposited in the NCBI Sequence Read Archive (SRA), yielding 6,100 unique sequencing runs after deduplication. Geolocated runs originated from 128 distinct cities across 19 countries. The sampling was concentrated in the United States (2,076 runs; mainly New York City, with additional sites such as Fairbanks and Denver), China (1,525 runs; mainly Shanghai and Shenzhen), and the United Kingdom (755 runs; mainly London).

**Figure 1.**
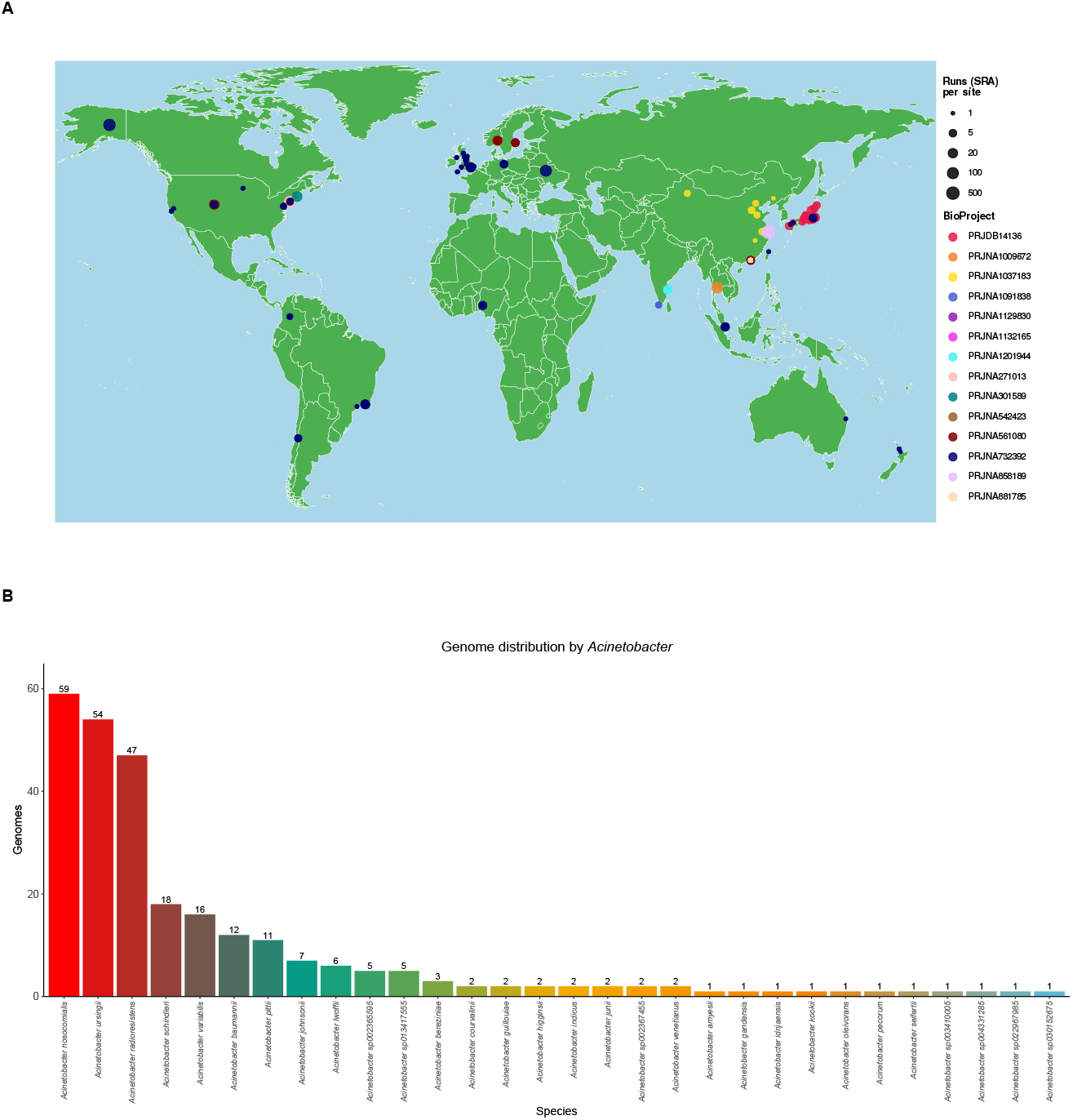
Projects processed and MAGs found. A) Map showing the geographic origin of the projects analysed. B) Acinetobacter species for which MAGs were found in the public transport system. The numbers on the top of the bars give the number of MAGs constructed for each species.

Further countries included Japan (319 runs; Tokyo, Narita, Fujisawa and other municipalities), Norway (242 runs; Oslo), Singapore (89 runs), Hong Kong (86 runs), Ukraine (86 runs; Kyiv), Brazil (71 runs; Rio de Janeiro, São Paulo), Sweden (64 runs; Stockholm), India (60 runs; Chennai, Thiruvananthapuram), Thailand (54 runs; Bangkok), Nigeria (42 runs; Offa, Ilorin), Chile (25 runs; Santiago), Germany (21 runs; Berlin), Colombia (18 runs; Bogotá), New Zealand (9 runs; Auckland, Hamilton), Taiwan (3 runs; Taipei) and Australia (2 runs; Brisbane). We noted that the isolation source was reported for 2,629 runs (43.1%); the most frequent sources were underground/subway environments, air, and other built-environment surfaces near subway stations. In total, we inferred 839 *Acinetobacter* MAGs, but only 269 MAGs had completeness > 95% and contamination < 5% and were considered for downstream analyses (see Supplementary Table 1). Most of the MAGs (98%, 263 out of the 269) came from subway systems (see Supplementary Table 1). Although we found MAGs for more than 20 well-defined *Acinetobacter* species, among them, *A. baumanii* (see Figure 1B), only 5 species had more than 15 MAGs; see the 5 bars in Figure 1B. The five species with the most MAGs were *A. nosocomilalis, A. radioresistens, A. schindleri, A. ursingii*, and *A. variabilis*. Thus, it seems that several species from the genus *Acinetobacter can* dwell in public transport systems; yet, only a few of them are the most prevalent. To have statistical power, downstream analyses were conducted for species having 15 or more MAGs, that is, the 5 species mentioned above.

### MAGs are an important source of both lineage diversity and gene diversity

To be able to properly appraise the diversity contributed by the MAGs, we included 685 genomes from RefSeq (see Methods and Supplementary Table 2). Then, core-genome phylogenies were constructed for each species (see Methods). In order to evaluate how much the MAGs contribute to lineage diversity, we computed the phylogenetic diversity (PD) for the trees considering only genomes and the trees including both MAGs and genomes. PD is a measure of diversity that is based on the phylogenetic relationships among the taxa considered and is implemented by adding all the branch lengths of the species tree [11]. As can be seen in Figure 2A and Table 1, for three species there is a clear increase in PD when MAGs are included. The most striking case is *A. variabilis*, where the inclusion of MAGs yielded a 91.2% increase in PD. Whereas for *A. radioresistens* and *A. schindleri*, the increases in PD were 48.71 and 38.29, respectively. Thus, for some *Acinetobacter* species, MAGs can contain a huge amount of diversity. Then, to analyse the diversity in terms of gene families, we also conducted pangenome analyses for the five species (see Table 2). In all the other species, we identified between 2100 and 3600 genes found only in the MAGs but not in the genomes. Clearly, these represent novel genes that have gone unnoticed with typical whole-genome sequencing. Remarkably, in some cases, the novel genes found in MAGs can account for more than 20% of the species’ total pangenome. For instance, *A. radioresistens* has 3685 (21.1%) gene families in MAGs out of the 17463 gene families in the whole pangenome; whereas *A. variabilis* has 2399 (23.03%) gene families uniquely in MAGs out of the total 10415 gene families. Collectively, these analyses show that MAGs represent not only understudied lineage diversity but also novel gene families. Of note, some of the 5 species have been barely studied, and actually no pangenome analyses are available for them. This is the case for *A. radioresistens, A. schindleri*, and *A. variabilis*; we conducted pangenome analyses for all three species, and our results show that they all have open pangenomes with large accessory genomes (see Figures 2B and 2C and Supplementary Figure 1). Furthermore, the three species had open pangenomes according to the heap’s law model. All in all, our results show that MAGs are a source of not only novel lineages but also many genes.

**Table 1.** Phylogenetic diversity analysis. PD for the trees containing just genomes as well as genomes and MAGs

| Species | PD genomes | PD genomes + MAGs | *PD increase |
| --- | --- | --- | --- |
| *A. nosocomialis* | 0.538884 | 0.570873 | 5.94 % |
| *A. ursingii* | 0.403911 | 0.44119 | 9.23 % |
| *A. radioresistens* | 0.210482 | 0.313015 | 48.71 % |
| *A. schindleri* | 0.322487 | 0.445974 | 38.29 % |
| *A. variabilis* | 0.094928 | 0.1815 | 91.2 % |

\*PD increase = 100\*(((PD genome + MAGs) – PD genomes)/PD genomes)

**Table 2.**
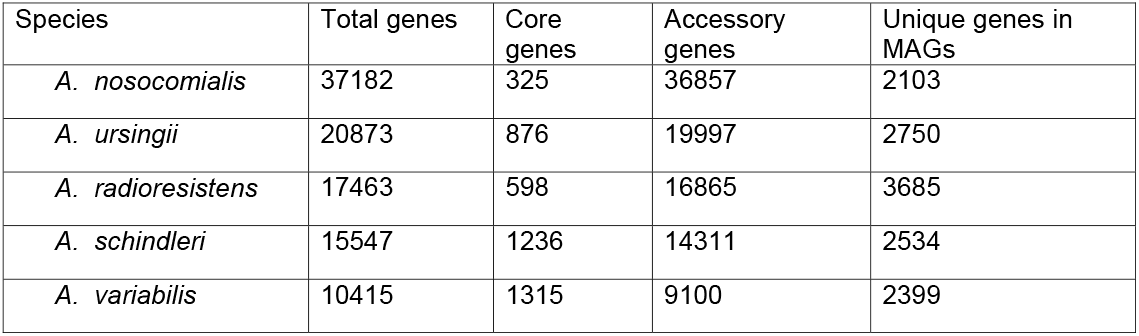
Pangenome analyses. Pangenome analyses of the 6 most relevant species

**Figure 2.**
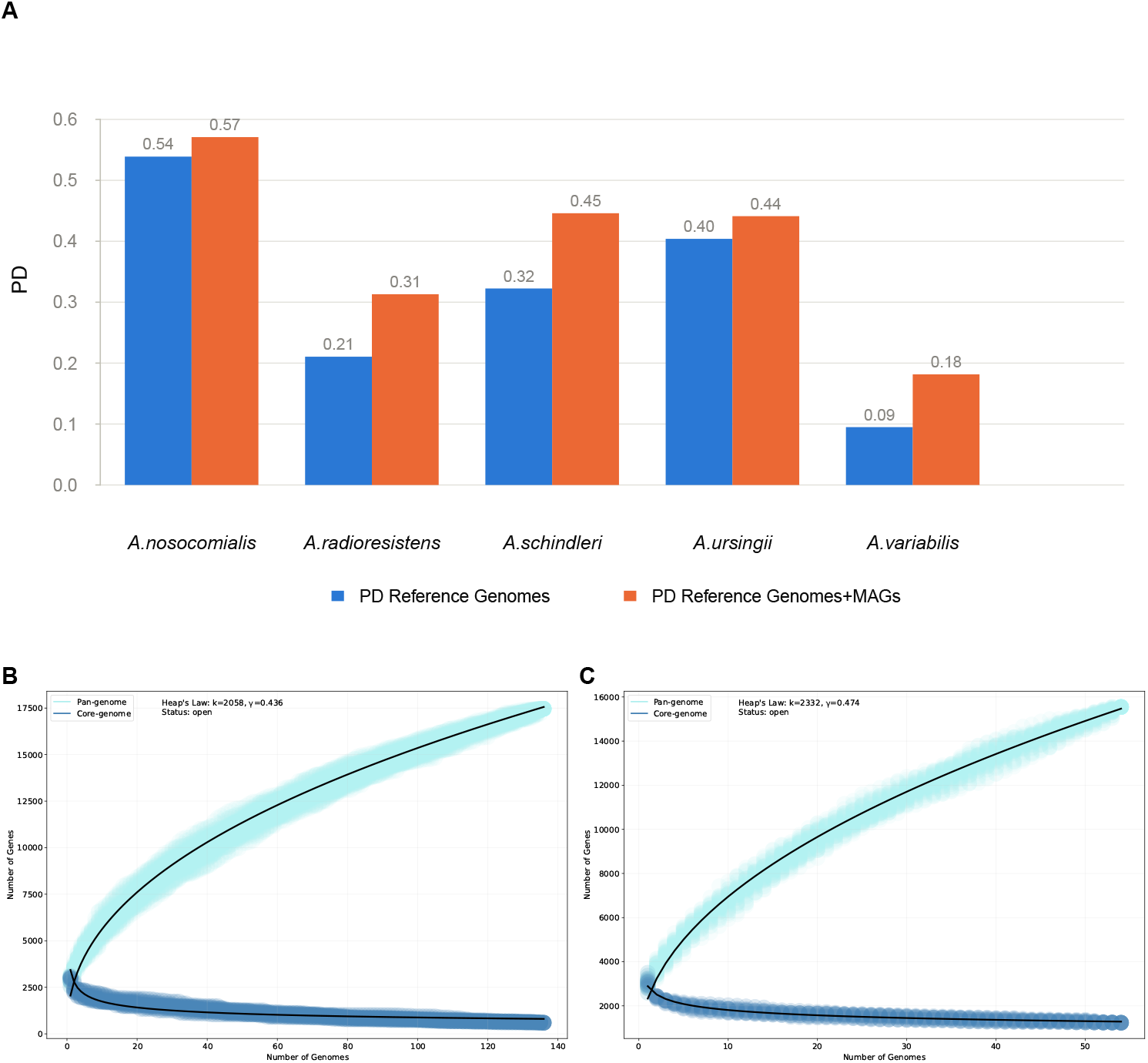
MAGs contain novel genomic diversity. A) Phylogenetic diversity for trees only containing genomes and trees containing both genomes and MAGs. B) Pangenome plot for *A. radioresistens;* light blue shading depicts how the pangenome increases as more genomes are added, whereas dark blue shading shows how the core genome decreases as more genomes are added. C) Pangenome plot for *A. schindleri*, light and dark blue shading the same as panel B.

### Genomic epidemiology of the MAGs

We then explore how the MAGs relate to the genomes recovered from different sources. Figure 3 and Supplementary Figure 2 show the phylogenies for the 5 species. In all the phylogenies, it is clear that MAGs did not group into one or a few clades, with one exception discussed below. Furthermore, the MAGs are located in diverse parts of the trees. Clear examples are the phylogenies of *A. nosocomilalis* (Figure 3A) and *A. radioresistens* (Figure 3B), where the MAGs are distributed across the trees and intermixed with clinical isolates, and sometimes even with animal isolates. These patterns also applied to *A. ursingii* (Figure 3D), *A. schindleri* (Figure 3C), and *A. variabilis*; although in the case of the last two, given that they contained fewer genomes and MAGs, the patterns are less salient. Importantly, for three species (*A. nosocomilalis, A. radioresistens*, and *A. ursingii*), we noted clear cases where MAGs and clinical genomes are very closely related, forming even polytomies in some instances. This indicates very recent transmission between hospitals and public transport settings. *A. ursingii* is the only case where we noted one rather big clade (bottom-right) composed exclusively of MAGs (Figure 3D). Then, to have a more precise idea of how the MAGs relate to isolates from clinical origin, we ran F_ST_ analyses for the 5 species. In this analysis, two groups were considered: MAGs on one side and genomes from human and clinical environments on the other. Of note, the fixation index (F_ST_) is a common measure of genetic differentiation [12], which can go from 0 to 1.

**Figure 3.**
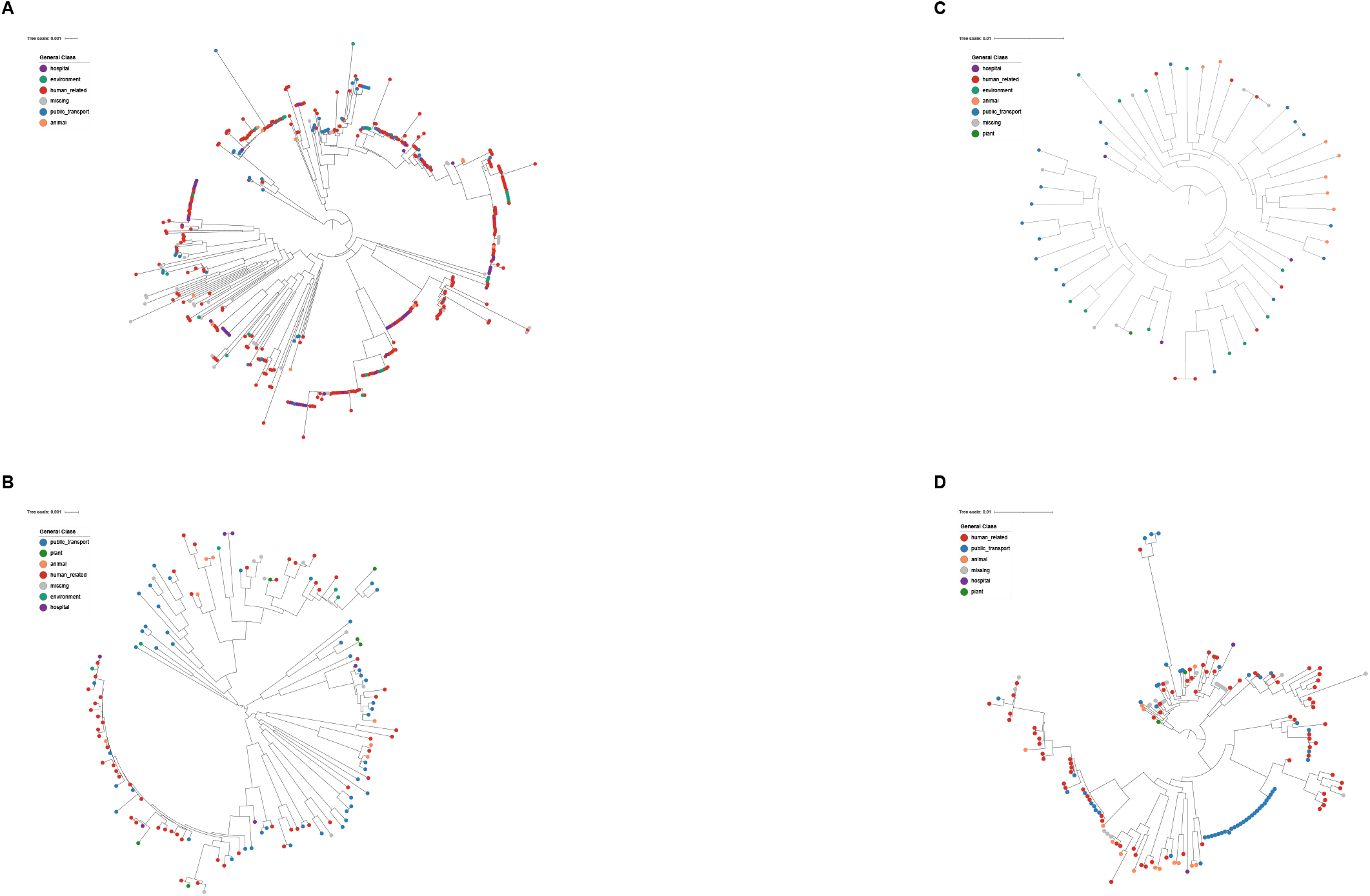
Core genome phylogenies. A) Core genome of *A. nosocomilalis*, B) Core genome of *A. radioresistens*, C) Core genome phylogeny of *A. schindleri*, D) Core genome of *A. ursingii*. In all four phylogenies, the scale bar gives the number of substitutions per site, and the source of the genomes and MAGs is color-coded; see color keys in each phylogeny.

Where 0 means little to no differentiation and 1 implies complete differentiation between populations. For four species, we noted very little differentiation between the MAGs and the genomes from clinical environments: *A. nosocomilalis*, (F_ST_ value = 0.048), *A. radioresistens* (F_ST_ value = 0.056), *A. schindleri* (F_ST_ value = 0.025), and *A. variabilis* (F_ST_ value = 0.039). The only species that showed a slightly moderate level of differentiation was *A. ursingii* with an F_ST_ value = 0.107. We think this higher value (compared with the other species) has to do with the big clade composed exclusively of MAGs discussed above. Collectively, the phylogenies and F_ST_ analyses show that the MAGs are not well differentiated from clinical genomes.

### Expanding the resistome of *Acinetobacter*

We then analysed the resistome of the public transport metagenomic samples of the five species. To do so, we implemented two approaches: one was identifying antibiotic resistance genes (ARGs) in the MAGs, but we also inferred the resistome for the whole metagenomic sample under study and then selected only ARGs belonging to the species under question using the reference genomes from such species (see Methods); we called the latter the *ad hoc* strategy. MAGs tend to recover considerably fewer ARGs than our *ad hoc* strategy (see Supplementary Figure 3). For instance, in *A. ursingii*, the *ad hoc* strategy identified 7-fold more ARGs than MAGs (see Figure 4A). Figure 4 and Supplementary Figure 3 provide the resistome for all five species. Considering the resistome of *A. radioresistens* (Figure 4B), we found very clinically important ARGs. For instance, we found several variants of the gene family *blaOXA*_*23*_, which codes for the carbapenemase OXA-23, the most widely acquired OXA in *A. baumannii*. We also noted the presence of sulfonamide-resistant dihydropteroate synthase genes *sul1* and *sul2*. Other important genes were the aminoglycoside nucleotidyltransferase *addA* and the tetracycline efflux pump gene *tet(39)*. In *A. schindleri* (Figure 4C), we found a couple of OXA-23 variants, along with 7 variants of another Oxacillinase (OXA-134). The genes *sul1, sul2, addA*, and *tet(39)* were also identified in this species. Notably, *A. nosocomilalis* was the only species with RND multidrug efflux system genes (see Supplementary Figure 4A). The species that had the fewest number of ARGs was *A. variabilis*, with 8 ARGs (see Figure 4D). Yet, among those ARGs, we found the clinically important genes *sul1* and *sul2*. Finally, we noted that some clinically relevant genes were found in most of the 5 species. The genes *sul1* and *sul2* were found in all five; *aadA* genes were found in four species, whereas *blaOXA*_*23*_ genes were found in three species. Then, to visualize how (dis)similar genomes from clinical origins are compared with MAGs, we ran Principal Coordinate Analyses on the ARG content (see Methods and Figure 5); we only did this for species with more than 10 different ARGs. We also evaluated whether the differences were statistically significant by conducting PERMANOVA analyses (see Figure 5). On the one hand (right-hand side, Figure 5), considering the ARG profiles of *A. radioresistens* and *A. schindleri*, MAGs and clinical genomes do not seem to cluster differently from one another; PERMANOVA p-values > 0.01. On the other hand (left-hand side, Figure 5), considering ARG profiles of *A. nosocomialis* and *A. ursingii*, MAGs and clinical genomes are more differentiated (PERMANOVA p-values < 0.01). In the case of *A. nosocomialis*, clear differentiation is noted along PCo1; whereas in the case of *A. ursingii*, notable differentiation is evident across PCo2.Collectively, our results show that MAGs underestimate the resistome; however, taking advantage of the reference genomes can significantly increase the estimation of ARGs when analysing metagenomic data. In addition, whereas for *A. nosocomialis* and *A. ursingii*, the ARG profiles of MAGs and clinical genomes are differentiated; in the case of *A. radioresistens* and *A. schindleri*, the ARG profiles of MAGs and clinical genomes are not statistically different. Finally, the five species have very important ARGs, such as *sul1, sul2, aadA, and blaOXA*_*23*_.

**Figure 4.**
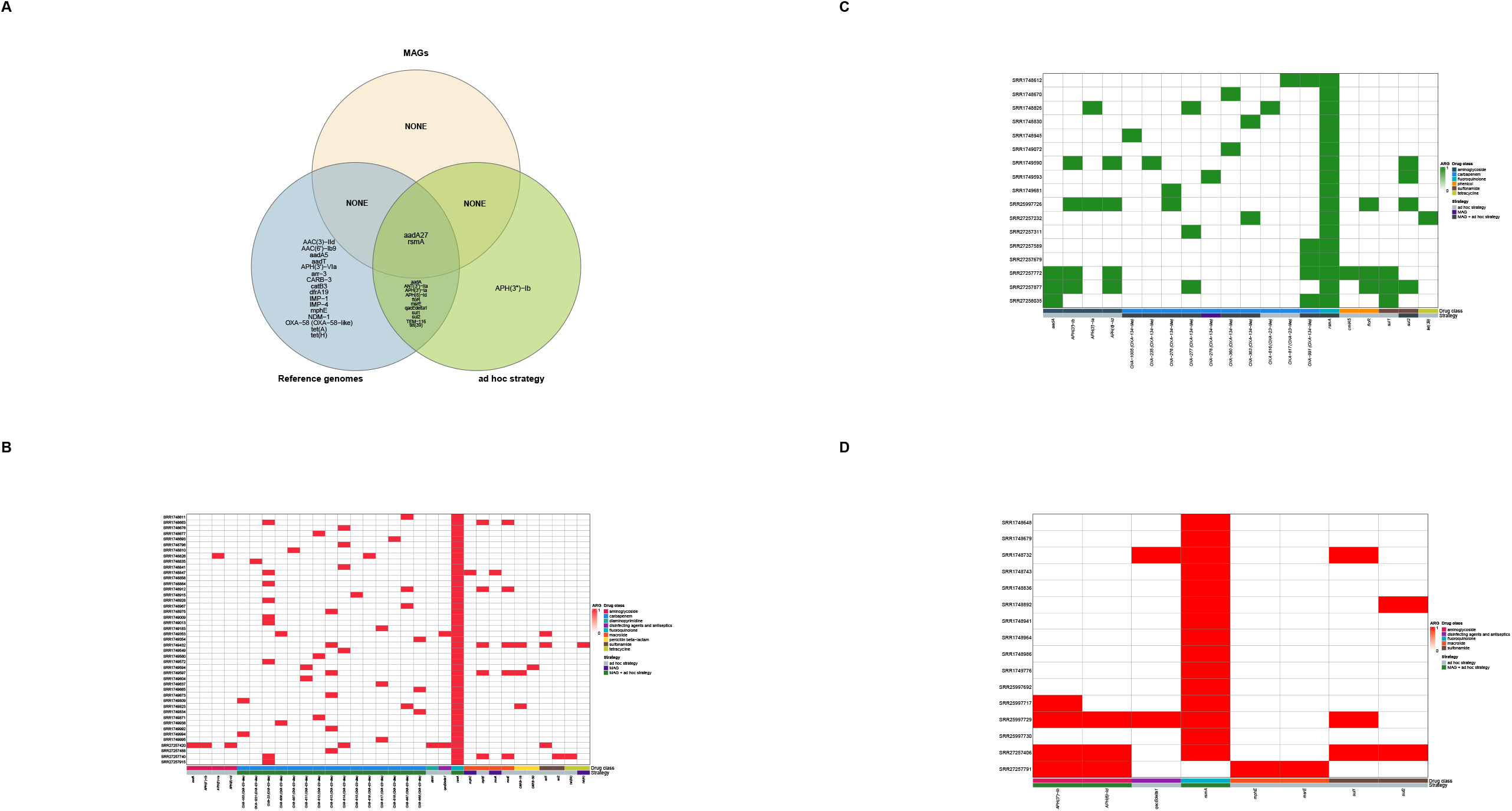
Resistome inference. A) Venn Diagram showing ARGs identified by MAGs and the *ad hoc* strategy in the case of *A. ursingii*. B) Heat map showing the resistome of *A. radioresistens*. C) Heat map showing the resistome of *A. schindleri*. D) Heat map showing the resistome of *A. variabilis*. For panels B to D, color vectors at the bottom show the drug class and the strategy that identified the genes.

**Figure 5.**
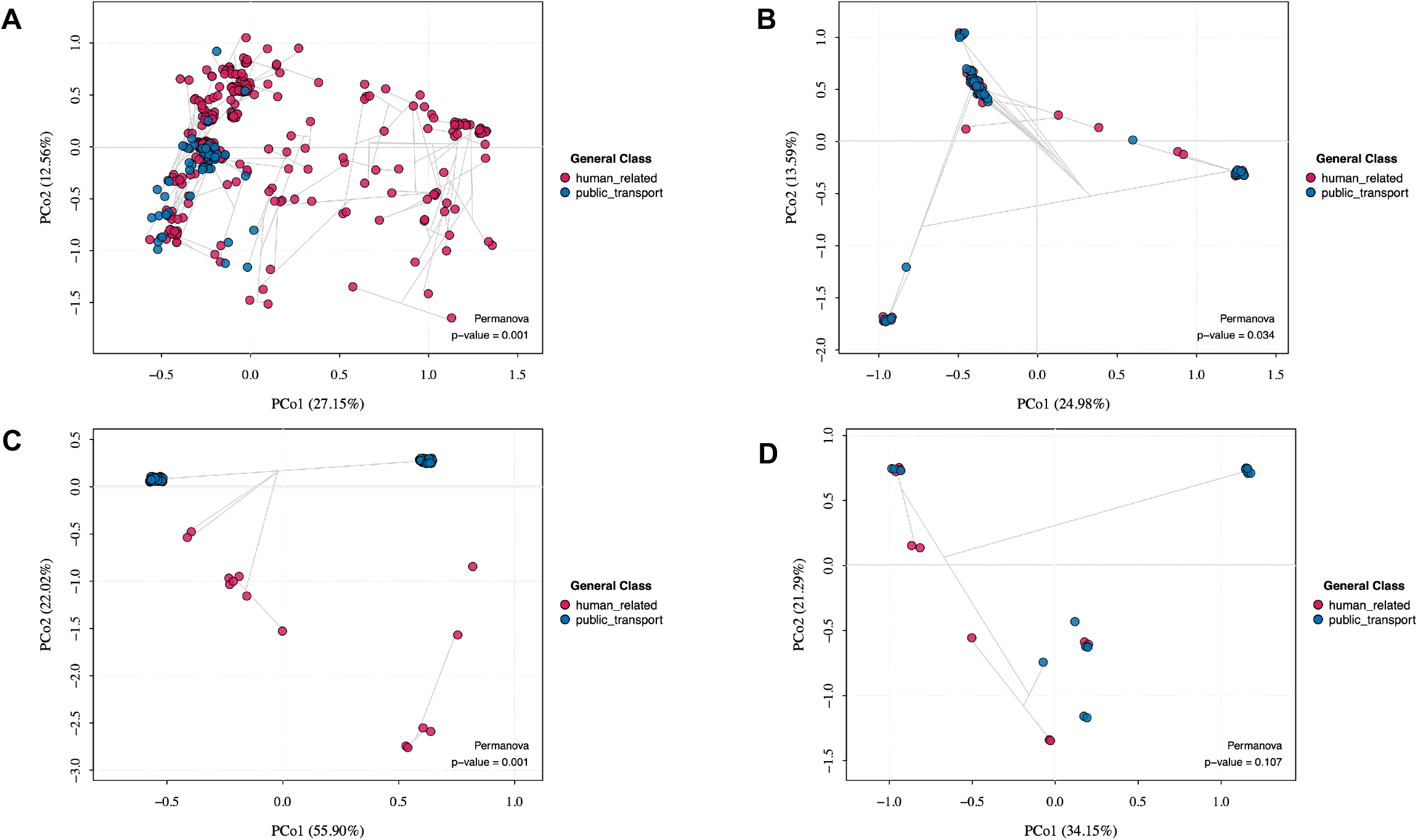
Resistome differentiation between clinical genomes and MAGs. A) PCoA and PERMANOVA analysis of *A. nosocomilalis* resistome, B) PCoA and PERMANOVA analysis of *A. radioresistens* resistome, C) PCoA and PERMANOVA analysis of *A. ursingii* resistome. D) PCoA and PERMANOVA analysis of *A. schindleri* resistome. In all four cases, blue dots (public_transport) denote the MAG, whereas red dots (human_related) show clinical isolates. Clinical isolates refer to human isolates but also isolates sampled from hospital environments.

## Discussion

The main idea of this study was to show that MAGs, by leveraging genomes, can be highly useful for understanding overlooked lineages and genes in *Acinetobacter*, especially from non-typical sources. We chose metagenomic data from public transport because public transport systems may play a role in disseminating different diseases. Our data suggest that several species, but mainly five, from the genus *Acinetobacter* are found in public transport settings. Remarkably, four of them (*A. ursingii, A. radioresistens, A. schindleri*, and *A. variabilis*) have been very little studied. MAGs from these species were a source of novel genetic diversity both at the lineage level and at the gene level. It is relevant to mention that the increase in PD and the number of novel genes (genes only located in MAGs) is likely to be correlated, to some extent, with the previous number of genomes sequenced. We noted that species that show the most conspicuous increases in PD and novel genes are those that have just a few genomes already sequenced. All in all, we think that MAGs are very useful for studying neglected diversity in those species with very little genomic information available.

Considering the genomic epidemiology of these MAGs, it is clear that for all five species, there is dissemination between the different settings. Both the phylogenies and the F_ST_ analyses show that the MAGs are not differentiated from clinical genomes. Moreover, in three species (*A. nosocomilalis, A. radioresistens*, and *A. ursingii*), there are cases of very recent transmission between public transport and clinical settings, where MAGs and clinical genomes are very closely related. The only conspicuous exception is just one clade in *A. ursingii*, which only has MAGs. These results agree with one of the scenarios that have been put forward for the dissemination of *A. buamannii*, where some population/lineages of *A. buamannii* are circulating across different sources or settings [4, 5]. Our genomic epidemiology results highlight a need for conducting more extensive surveillance where we pay as much attention to the non-human, non-clinical settings as we do to the clinical settings [13, 14].

In terms of the resistome, although we did not find as many ARGs as those detected in *A. buamannii* [15], we found some very important ARGs from a clinical point of view. For instance, *sul1* and *sul2* were found in the five species, whereas *aadA* was found in four. Other relevant ARGs were *blaOXA*_*23*_ and *tet(39)* that were found in 3 of the species. From a methodological point of view, we want to highlight that MAGs are clearly underestimating the number of ARGs. To amend that, we developed an *ad hoc* strategy that helps us to recover more ARGs. Our data suggest that the *ad hoc* strategy recovers considerably more ARGs than the MAGs. For instance, in the case of *A. ursingii*, the *ad hoc* strategy identified 7-fold as many ARGs as those found in MAGs. Yet, it is important to mention that some ARGs could only be extracted via MAGs; thus, in light of this, we proposed that using both MAGs and our *ad hoc* approach is the best way to get a more reliable estimate of the resistome for the lineages inhabiting public transport. Finally, in terms of how similar the resistome of the clinical genomes is to the MAGs, we noted that whereas *A. nosocomilalis* and *A. ursingii* show differentiation, that was not the case for *A. radioresistens* and *A. schindleri*, where the PCoA and PERMANOVA analyses do not tell apart the MAGs from the clinical genomes.

In conclusion, our results show that MAGs can be a source of novel diversity both at the lineage and gene level. Also, MAGs from non-standard settings help to explain the transmission dynamics of *Acinetobacter* species outside of the clinic and between different settings. Yet, as far as the resistome is concerned, in addition to considering MAGS, *ad hoc* strategies (leveraging reference genomes) are needed to more accurately predict the actual number of ARGs.

## Methods

### Metagenomic and genomic data retrieval and sample selection

Public transport metagenomic datasets were retrieved from the National Center for Biotechnology Information (NCBI). Geographic metadata were standardized using tidyverse [16] and geocoded with tidygeocoder [17] through OpenStreetMap/Nominatim. Sampling localities were visualized on global maps using ggplot2 and maps [18]. Each dataset was taxonomically classified using Kraken2 v-2.1.3 [19] with the MiniKraken2 reference database. Based on prior empirical observations from successfully reconstructed *Acinetobacter* MAGs, only datasets containing at least 43,000 reads taxonomically assigned to the genus *Acinetobacter* were retained for further exploration. Low-quality sequences were removed using Trimmomatic v-0.39 [20], applying a minimum Phred quality score threshold of Q30. All the *A. nosocomialis, A*.*ursingii, A*.*radioresistens, A*.*schindleri*, and *A. variabilis* genomes were downloaded from REFSEQ and GenBank, excluding Metagenomic Assembled Genomes (MAGs), as of February 16, 2026. We included genomes that had completeness ≥ 95% and contamination ≤ 5%; both measures were determined by running CheckM v1.2.2 [21] on the genomes. We made sure that the genomes were correctly assigned to *Acinetobacter* species by conducting an average nucleotide identity (ANI) analysis for all the genomes using GTDB-Tk v2.4 [22], which requires an ANI value ≥ 95% to the reference species. Metadata was downloaded from the NCBI https://www.ncbi.nlm.nih.gov/datasets/genome/ webpage. In Supplementary Table 1 are all the MAGs, and genomes along with the accession numbers of the metagenomic BioProjects and metadata (in the case of MAGs) lists all these genomes with their accession numbers and metadata.

### Metagenomic assembly, genome binning and refinement

Filtered metagenomic reads were assembled *de novo* using SPAdes v4.0.0. [23]. To improve assembly quality and downstream binning performance, contigs shorter than 1.5 kb were removed using SeqKit v2.10.0 [24], retaining only contigs ≥1.5 kb for subsequent analyses. Contig coverage information was generated using the gen_cov_file.sh script provided by COMEBin v1.0.4 [25]. Genome binning was performed independently using MetaDecoder v1.2.1 [26], SemiBin2 v 2.2.0 [27], and COMEBin v1.0.4. To improve bin quality and consensus, binning outputs from all three tools were integrated and refined using DAS Tool v1.1.7 [28].

### Taxonomic classification, quality assessment, pangenome analysis, and phylogenetic reconstruction

Refined bins were taxonomically classified using GTDB-Tk v2.4.1[9] to identify MAGs corresponding to the *Acinetobacter* genus. Bin completeness and contamination were assessed using CheckM v 1.2.2 [21]. Only MAGs with completeness of ≥ 95% and contamination ≤ 5% were retained for downstream analyses. MAGs exceeding 5% contamination were further processed using MAGPurify2 v1.0.2 [29] to remove duplicated or contaminant contigs, when possible, and were re-evaluated after purification. MAG annotation for the species *A. nosocomialis, A*.*ursingii, A*.*radioresistens, A*.*schindleri*, and *A. variabilis* was conducted using Prokka v 1.14.6 [30]. MAGs from those species were included in a pangenome analysis together with additional genomes downloaded from REFSEQ and GENBANK using Roary v3.13.0 [31]. Single-copy gene family genes identified from the pangenome were extracted and used to construct a phylogenetic tree, enabling comparative genomic and evolutionary analyses. Maximum-likelihood (ML) phylogenies were generated with RAxML [32] and processed using ape [33]. Maximum-likelihood phylogenetic trees were inferred employing the GTR nucleotide substitution model with invariant sites and gamma-distributed rate heterogeneity GTR+I+G4. Faith’s phylogenetic diversity (PD) was estimated with picante [34] from the maximum-likelihood phylogenies generated. Phylogenetic trees were visualized and plotted using the iTOL v7 tool [35].

### Resistome profiling

Antimicrobial resistance genes were identified by screening *Acinetobacter* MAGs against the Comprehensive Antibiotic Resistance Database (CARD) database v-3.2 [36]. In this analysis, an identity percentage of ≥60% and a coverage of ≥70% were considered for determining the antibiotic resistance genes (ARGs). The presence and distribution of resistance determinants across MAGs were subsequently analyzed. In the *ad hoc* strategy, the resistome for the whole metagenome (sample analysed) was conducted, also with an identity percentage of ≥60% and a coverage of ≥70%, and then ARGs for the species under question were filtered as follows. To identify homologous ARGs associated with each of the five species, we conducted BLAST searches of the genes from the reference genomes against the resistome for the whole metagenome. For the BLAST searches, we used BLASTn v-2.17 [37], with an E-value of 1e-20, a minimum sequence identity of 95%, and a minimum alignment coverage of 90%. BLAST homology criteria were established to identify homologous gene sequences at the species level. Binary ARG profiles were visualized using ComplexHeatmap [38], with genes grouped according to drug class. Resistome composition was assessed using binary Jaccard dissimilarities, principal coordinates analysis (PCoA), and PERMANOVA implemented in vegan [39, 40]. All statistical analyses and visualizations were performed in R [41].

## Data Sharing

Supplementary Table 1 lists all the MAGs constructed in this study, whereas Supplementary Table 2 lists all the genomes used in this study. These tables provide the GenBank or RefSeq assembly accession numbers for all the sequences.

## Authors contributions

SCR conceived and coordinated the project. RLS gathered the data. AAV and RLS curated the data collection. AAV developed the pipelines for constructing the phylogenies and predicting the resistome. RLS performed the bioinformatic analyses. SCR drafted the article, and all the authors approved the final version of the article.

## SUPPLEMENTARY FIGURES

**Supplementary figure 1**

Pangenome plot for *A. variabilis*, with light and dark blue shading the same as Figure 2B.

**Supplementary figure 2**

Core genome phylogeny of *A. variabilis*.

**Supplementary figure 3**

Comparison of AGRs found by MAGs and the *ad hoc* strategy. Comparisons were carried out for the 5 species. Asterisks denote statistically significant differences.

**Supplementary figure 4**

Resistome of *A. nosocomialis* and *A. ursingii*.

A) Heat map showing the resistome of *A. nosocomialis*; B) Heat map showing the resistome of *A. ursingii*. Color vectors at the bottom show the drug class and the strategy that identified the genes.

## SUPPLEMENTARY TABLES

**Supplementary Table 1**

List of all the MAGs constructed in this study. Completeness, contamination, BioProject, SRA, BioSample numbers, and other metadata are provided for each MAG.

**Supplementary Table 2**

List of all the genomes used in this study. Metadata as well as the accession numbers are provided for all the genomes considered.

