## Supplementary figures and images for "Leveraging genomes and metagenome-assembled genomes unveils novel diversity and transmission between public transport and clinical settings in *Acinetobacter*"

### Supplementary Figure 1

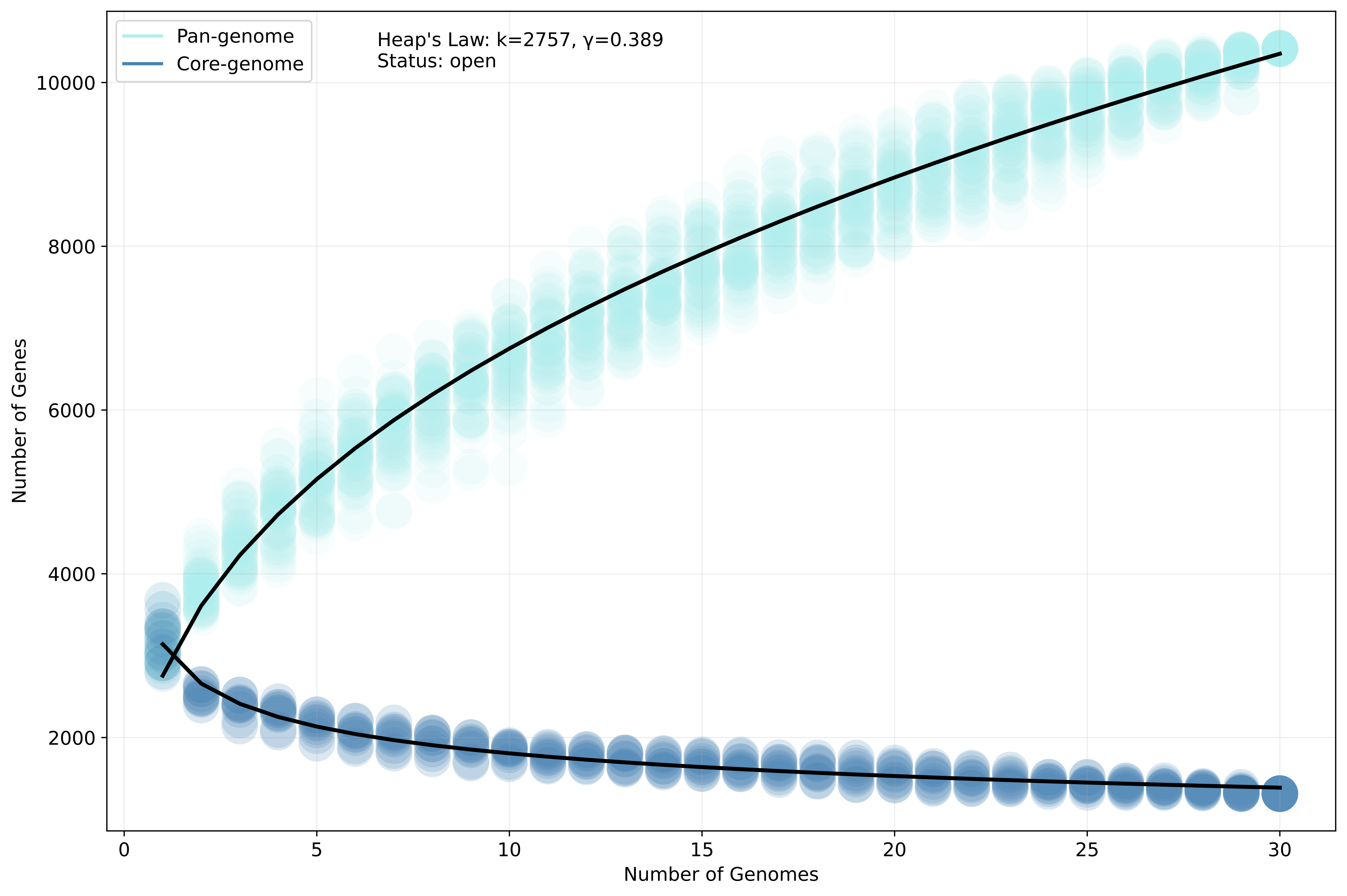

### Supplementary Figure 2

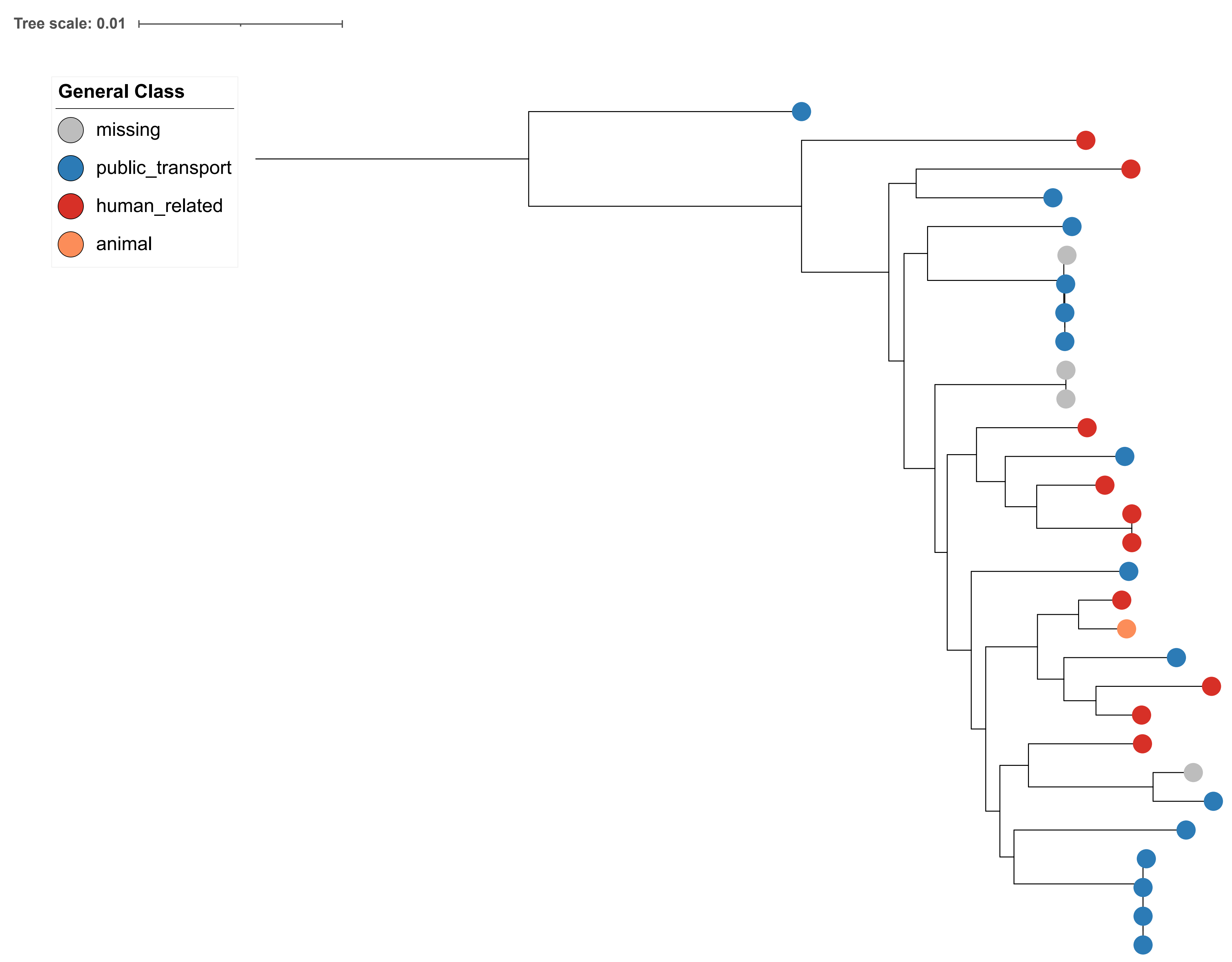

### Supplementary Figure 3

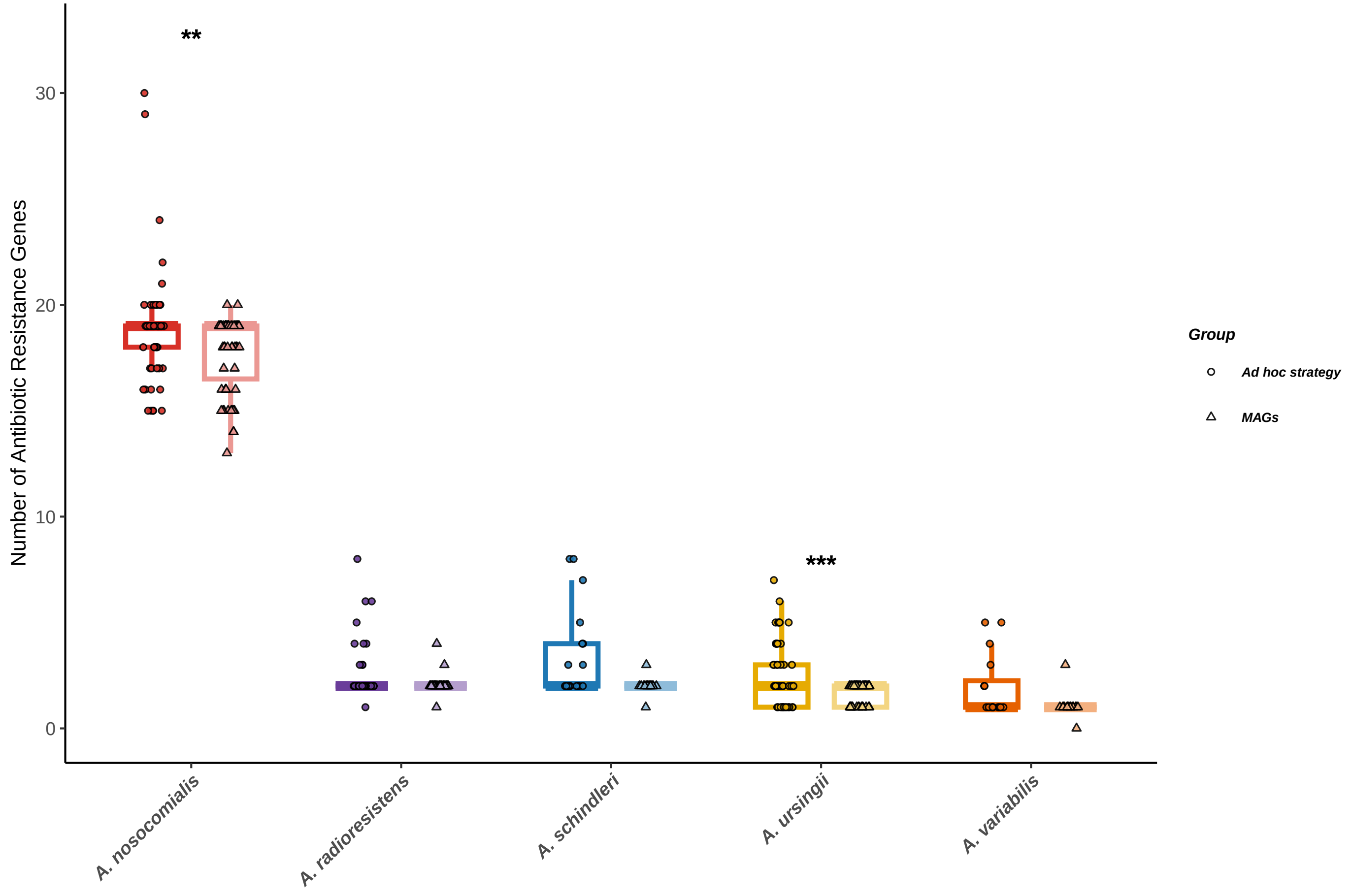

### Supplementary Figure 4

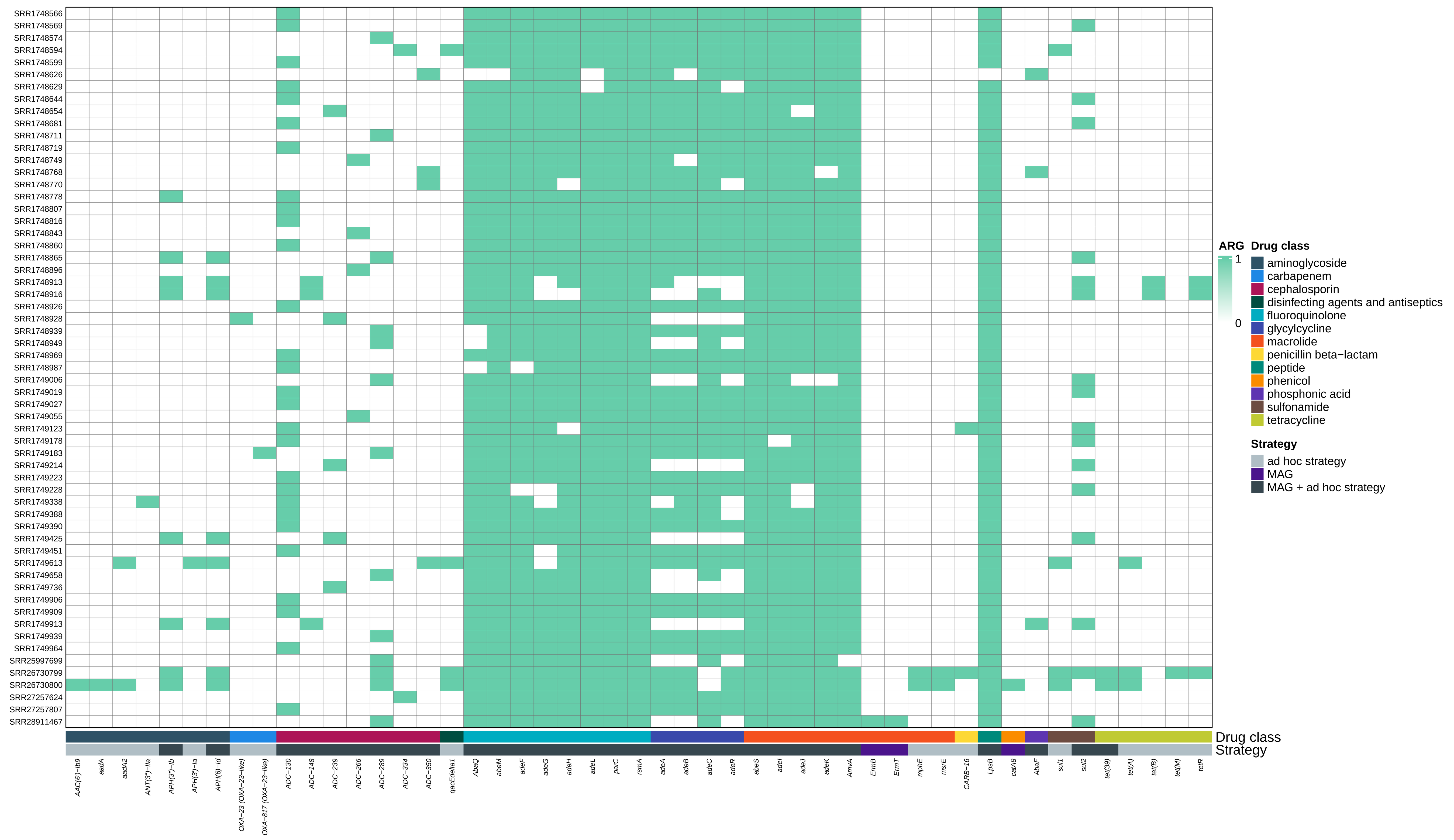

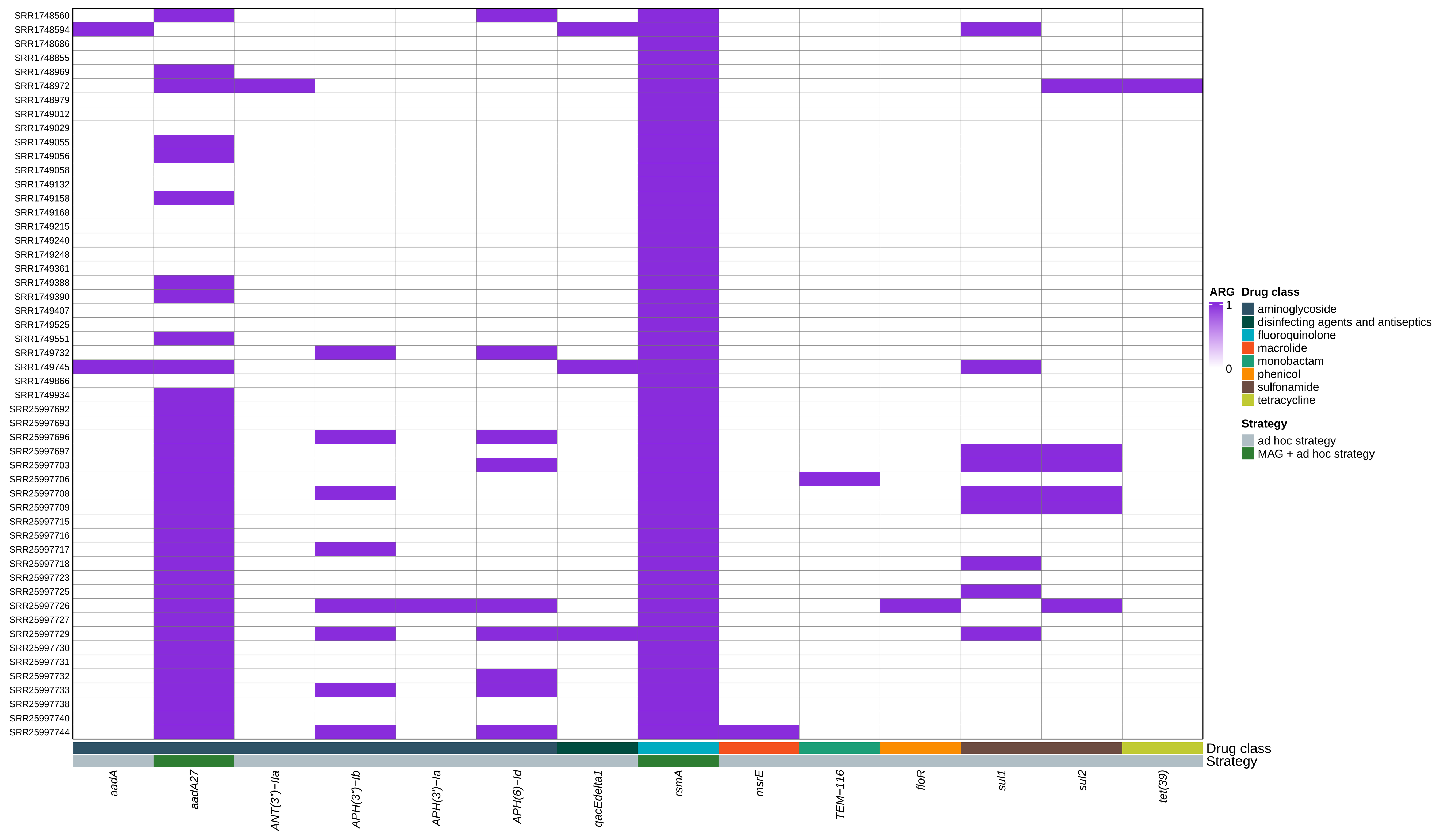
